# Phosphorylation of Serine114 of the transcription factor ABSCISIC ACID INSENSITIVE 4 is essential for activity

**DOI:** 10.1101/2020.08.14.250936

**Authors:** Nadav Eisner, Tzofia Maymon, Ester Cancho Sanchez, Dana Bar-Zvi, Sagie Brodsky, Ruth Finkelstein, Dudy Bar-Zvi

## Abstract

The transcription factor ABA-INSENSITIVE(ABI)4 has diverse roles in regulating plant growth, including inhibiting germination and reserve mobilization in response to ABA and high salinity, inhibiting seedling growth in response to high sugars, inhibiting lateral root growth, and repressing light-induced gene expression. ABI4 activity is regulated at multiple levels, including gene expression, protein stability, and activation by phosphorylation. Although ABI4 can be phosphorylated at multiple residues by MAPKs, we found that S114 is the preferred site of MPK3. To examine the possible biological role of S114 phosphorylation, we transformed *abi4-1* mutant plants with *ABI4pro::ABI4* constructs encoding wild type (114S), phosphorylation-null (S114A) or phosphomimetic (S114E) forms of ABI4. Phosphorylation of S114 is necessary for the response to ABA, glucose, salt stress, and lateral root development, where the *abi4* phenotype could be complemented by expressing ABI4(114S) or ABI4(S114E) but not ABI4(S114A). Comparison of root transcriptomes in ABA-treated roots of *abi4-1* mutant plants transformed with constructs encoding the different phosphorylation-forms of S114 of ABI4 revealed that 85% of the ABI4-regulated genes whose expression pattern could be restored by expressing ABI4(114S) are down-regulated by ABI4. Over half of the ABI4-modulated genes were independent of the phosphorylation state of ABI4; these are enriched for stress responses. Phosphorylation of S114 was required for regulation of 35% of repressed genes, but only 17% of induced genes. The genes whose repression requires the phosphorylation of S114 are mainly involved in embryo and seedling development, growth and differentiation, and regulation of gene expression.

**Highlights:**

- Transcription factor ABI4 is a substrate of MAP kinases.
- MPK3 preferentially phosphorylates Serine 114 of ABI4.
- Phosphorylated Serine 114 of ABI4 is required for the complementation of abi4 mutants.
- Phosphorylated ABI4 acts primarily as a repressor.

## 1. Introduction

The transcription factor ABA-INSENSITIVE(ABI)4 was initially identified on the basis of ABA-resistant germination in loss of function mutants [1, 2]. Additional mutants were found in screens for resistance to stress responses due to high salinity [3], high sugar concentrations [4–6] or inhibited sugar-induced gene expression [7]. Further characterization of these mutants revealed roles in lipid reserve mobilization, lateral root growth, regulation of light-regulated gene expression, retrograde signaling from mitochondria, redox signaling, and some pathogen responses (reviewed in [8]). ABI4 has also been implicated in retrograde signaling from plastids [9], but this is controversial [10].

Although *ABI4* is most strongly expressed in embryos, imbibing seeds and maturing pollen [11–13], its diversity of functional roles reflects expression in vegetative tissues. In addition to developmental contexts, *ABI4* promoter activity is regulated by environmental stresses and interactions with signaling via other hormones including cytokinin and auxin [14–16]. ABI4 protein accumulation is further regulated by proteasomal degradation, but is stabilized in the dark or by exposure to high salt or sugar concentrations [17–19].

Like many cellular proteins, ABI4 protein activity may also be subject to regulation by post-translational modifications such as phosphorylation. The primary amino acid sequence of ABI4 is highly enriched in the phosphorylatable residues Ser and Thr: 47 and 20 residues, respectively, out of the 328 amino acid residues, corresponding to 20% of the total amino acid residues of ABI4. An *in vitro* phosphorylation study of the Arabidopsis proteome by MAPKs showed that ABI4 could be phosphorylated by 3 out of the 10 tested MAPKs [20]. The phosphorylating MAPKs (MPK3, MPK4, and MPK6) regulate numerous signaling pathways, including stress response, in Arabidopsis [21, 22] and their activities were shown to be activated by ABA treatment, suggesting that this could be a mechanism for stress-regulation of ABI4 activity. MAPK signaling may also regulate ABI4 activity at the level of expression via effects on as-yet unidentified transcription factor(s) [23].

Previous studies of ABI4 phosphorylation focused on its role in retrograde signaling controlling light-regulated gene expression [24] and inhibiting adventitious root formation [25]. Guo et al. [24] demonstrated Ca^2+^-dependent scaffolding of MAPK cascades resulting in activation of MPK3 and MPK6, which in turn activated ABI4 to repress the photosynthesis-associated nuclear gene LHCB. In the present study, we tested the importance of ABI4 phosphorylation by MAPKs in regulating ABA and stress responses in seeds and seedlings.

## 2. Materials and Methods

### 2.1. Expression and purification of recombinant proteins

cDNA sequences of Arabidopsis genes encoding ABI4, MPK3, MPK4, and MPK6 were amplified by a polymerase chain reaction (PCR) using gene-specific primers. Resulting sequences were subcloned into the *SalI* and *XhoI* sites of pGST-parallel1 bacterial expression vector. The resulting plasmids were verified by DNA sequencing and introduced into *E. coli* BL-21 cells. The respective GST-MAPK proteins were expressed and affinity-purified on GST-agarose column.

The coding sequence of ABI4 was cloned into the *Xho*I and *Sal*I sites of pRSET A, downstream to a sequence encoding hexahistidine [26]. Site-directed mutagenesis was carried out according to the QuikChange protocol (Agilent Technologies). The primers used for mutagenesis are shown in Supplemental Table S1 using the pRSET A-ABI4 plasmid as a template. The introduced respective mutations were confirmed by DNA sequencing. Plasmids were introduced into *E. coli* BL-21 cells and cultures were grown at 16 °C in LB medium containing in addition 100 mg/ml ampicillin, to OD600 nm=0.6. The expression of the recombinant protein was induced by the addition of 1 mM IPTG. The culture was incubated overnight in the shaker at 16 °C. Bacteria were then harvested, and cells were osmotically shocked to reduce the abundance of cation chelators within the bacterial intermembrane space, which was shown to reduce the binding capacity of low-expressed recombinant His-tagged proteins to matrices containing immobilized nickel ions [27]. Hypertonically shocked bacteria were suspended in a lysis buffer containing 50 mM Tris-HCl pH 8.0, 0.5 M NaCl, and 10 mM imidazole, using a ratio of 1.5 buffer ml/gr bacteria and sonicated on ice by 60 seconds of 70Ω bursts using a tip sonicator (Sonics Vibra-Cell). The resulting homogenate was centrifuged for 45 min at 150,000 × g and 4 °C in an ultracentrifuge, and the resulting supernatant was loaded on a 1ml HiTrap IMAC FF nickel-agarose column (GE) attached to an AKTA start chromatography apparatus. The column was thoroughly washed with the lysis buffer and then eluted with a 20–300 mM imidazole gradient in a buffer containing 50 mM Tris-HCl pH 8.0 and 0.5 M NaCl. Eluted fractions were analyzed by SDS-PAGE; fractions containing ABI4 were pooled, and their buffer was replaced by 50 mM Tris-HCl pH 8.0 and 0.5 M NaCl by using a 10 kDa cutoff Amicon Ultra 0.5ml centrifugal filter (Merck).

### 2.2. Protein phosphorylation assay

Approximately 0.5 μg of affinity-purified His-tagged ABI4 proteins (WT and proteins containing the respective amino acid changes) were incubated in a phosphorylation reaction mix containing 20 mM HEPES-NaOH pH 7.4, 15 mM MgCl_2_, 5 mM EDTA, 1 mM DTT, 3 μl of carrier-free [γ-^32^P]ATP (3000 Ci/mmol; 5 mCi/ml), and 0.1μg of each of recombinant GST-MAPK3, GST-MAPK4, or GST-MAPK6 proteins (in phosphate-buffered saline, PBS). Reaction mixtures were incubated for 1 hour at 30 °C, followed by adding 10 μl of 5 × SDS-PAGE sample buffer. Proteins were denatured by incubation for 5 min at 95 °C, and samples were loaded on SDS-PAGE [28]. Following the electrophoresis run, the gel was stained with Coomassie blue, vacuum dried for 2 hours at 80 °C, and exposed to a phosphorimager screen. Signals were detected by a Typhoon phosphorimager (GE Life Sciences).

### 2.3. Plant material and growth

All the studies used *Arabidopsis thaliana*–Columbia ecotype. Seeds of *abi4-1* mutants were obtained from an Arabidopsis stock center in Columbus, Ohio. The plants were grown at 22–25 °C in day/night periods of 12 hours each and 50% humidity. For seed collection, plants were grown in long daylight regimes (16/8 hours, day/night) to accelerate flowering time. Seeds were surface sterilized by incubation for 20 min with constant rotation in 20% commercial bleach containing 1% Triton X-100, followed by eight washes with sterile deionized water. Seeds were resuspended in soft agar and incubated for 3–5 days at 4 °C, to enhance germination rate and uniformity. Cold stratified seeds were sown in Petri dishes containing solidified 0.5 × Murashige and Skoog (MS, [29]), 0.5% sucrose and 0.55% plants agar, containing, also, the added materials as indicated. Plants were also sown in 6 × 6 × 7 cm black pots using a potting mix containing 20 % tuff, 30% peat, 30% vermiculite, 20% perlite, and slow-release fertilizer. Pot-flats were placed in trays and were irrigated by filling the trays with deionized water.

### 2.4. Preparation of constructs and transformation of plants

DNA sequence comprised of ABI4 encoding sequence and 2 kbp of upstream promoter sequence was cloned by PCR using gene-specific primers (Supplemental Table S1) and template comprised of genomic DNA prepared form WT Arabidopsis plants. The resulting PCR product was verified by DNA sequencing and subcloned into the *Kpn*I and *Xba*I sites of the plant transformation vector pCAMBIA 2300. Constructs expressing ABI4(S114A), ABI4(S114E), ABI4(S130A) or ABI4(S130E) mutated proteins were constructed by replacing the *BamH*I-*Blp*I restriction fragment of the wild-type ABI4 in pCAMBIA 2300 by the respective fragments of constructs in pRSET encoding the respective mutation. The resulting plasmids were verified by DNA sequencing and were introduced into *Agrobacterium tumefaciens* (strain GV3101). The transformed bacteria were used to transform *abi4* mutant Arabidopsis plants [30]. Transgenic plants were selected by germination in Kanamycin containing plates. Homozygous seeds from the T2 and T3 generations, resulting from a single T-DNA insertion event, were chosen for this study.

### 2.5. Plant phenotyping

#### 2.5.1. Plate grown seedlings

Cold-treated, surface-sterilized Arabidopsis seeds were plated on Petri dishes containing 0.5 x MS, 0.5% sucrose, and 0.55% plant agar, supplemented with the indicated concentrations of ABA, glucose or NaCl. Plates were incubated in the growth room in 12 h of dark/light conditions at 22 °C, and radicle emergence was assayed daily for one week. Cotyledon development was assayed by measuring seedlings with spread cotyledons. Lateral root development was assayed in 12 days old seedlings grown on Petri dishes containing 0.5 × MS, 0.5% sucrose, and 0.5% plant agar. Seedlings were harvested, placed on a transparent sheet, and roots were arranged before taking their images by a scanner. The lengths of primary and lateral roots were measured from the resulting images. All assays were carried out at least 3 times with more than 50 seedlings per line in each assay. Statistical significance of the data was analyzed using Tukey’s HSD post-hoc test.

#### 2.5.2. Salt treatment of pot-grown plants

Seeds of plants of the indicated genotypes were sowed in 6 × 6 × 7 cm pots. The pots were placed in the growth room under conditions of 12h of dark/light at 22 °C for approximately 1 month. Plants were then irrigated with water or with a solution containing 0.25 M NaCl and 16 mM CaCl_2_. Plants were followed daily for 3 weeks. Rosettes having even a small amount of green tissue were scored as live plants, where all yellow-brown rosettes were scored as dead plants. All assays were carried out at least 3 times with more than 15 seedlings per line in each assay. Statistical significance of the data was analyzed using Tukey’s HSD post-hoc test.

### 2.6. RNA Sequencing and expression analysis

Arabidopsis seeds were plated on top of a sterilized silk print mesh placed on growth medium (0.5 x MS, 0.5% sucrose and 0.5% agar) and grown for 1 week as described above. Nets with the germinated seedlings were transferred to fresh Petri dishes containing Whatman No 1 filter paper soaked with 0.5 x MS, 0.5% sucrose and 50 μM of the plant hormone ABA. Plates were incubated in the light at room temperature for 4 h, roots were harvested, and total RNA was extracted by using Quick-RNA Plant Kit (Zymo Research). The RNA concentrations were determined by using NanoDrop (DeNovix DS-11 FX). Four biological repeats, each containing all tested genotypes, were performed on different days.

Sequencing libraries were prepared and sequenced by the Israel National Center for Personalized Medicine (INCPM) at the Weizmann Institute. Briefly, total RNA was fragmented, followed by reverse transcription and second-strand cDNA synthesis. The double-strand cDNA was subjected to end repair, A base addition, adapter ligation, and PCR amplification to create libraries. Libraries were evaluated by Qubit and TapeStation. Sequencing libraries were constructed with barcodes to allow multiplexing of the samples on two lanes of Illumina HiSeq 2500 V4 instrument, resulting in ~25 million single-end 60-bp reads per sample. Poly-A/T stretches and Illumina adapters were trimmed from the reads using cutadapt [31]; resulting reads shorter than 30 bp were discarded. Reads were mapped to the *A. thaliana* reference genome TAIR10 using STAR [32], supplied with gene annotations downloaded from Ensembl (and with EndToEnd option and outFilterMismatchNoverLmax was set to 0.04). Expression levels for each gene were quantified using htseq-count [33], using the annotation of the reference genome above.

The total reads of each sample were normalized to 20 million, and log2 transformed. Values below 2 (log2) were removed, and only genes where at least two out of the 18 samples had a threshold value of 6 (log2) were included in the analysis. Data were then normalized to remove the day bias: for each gene, the median value of all samples from the same day was subtracted from the value of each sample. Three outlier samples were not included in the analysis.

In order to find the genes regulated by the WT, we first compared it to *abi4*. In each comparison, genes were assigned to 30 bins according to the sum of expression of the two genotypes (values used for binning are log2 values prior to day normalization, for indication of absolute expression levels). Z-score of the delta log2 expression between the two genotypes was calculated for each bin, using the values normalized by experiment day. A cutoff of |Z-score| > 2 was implemented in order to find genes that are differentially expressed. The same was done in order to find genes that are differentially expressed between the *abi4* mutant and the transgenic lines expressing ABI4(114S), ABI4(S114A) or ABI4(S114E). Since expressing wild-type ABI4(114S) in the *abi4* genetic background complemented the wild-type phenotype (Figures 3-6), we focused our analysis on genes whose expression could be complemented by ABI4(114S). In order to find the genes that are affected by the specific phosphorylation in S114, we checked which of those genes could also be complemented by expressing the phosphorylation-null ABI4(S114A) or the phosphomimetic ABI4(S114E) in *abi4*.

ABI4 DNA binding sites were searched by using the Patmatch tool at TAIR. DNA sites searched were: S box (CACYKSCA), CE-1 like (CACCK), ABRE (YACGTGGC), ABRE-like (BACGTGKM), DRE core motif (RCCGAC) [34], and ABE motif (GCSCTTT) [26]. The resulting list was searched for genes whose expression in *abi4* mutants compare with wild-type plants, could be complemented in the *abi4* plants expressing ABI4(114S), ABI4(S114A) or ABI4(S114E). As a control we used the list of genes whose expression was detected by RNA-seq.

Gene Ontology (GO) was performed using the agriGO v2.0 (http://systemsbiology.cau.edu.cn/agriGOv2/) [35] using the Fisher statistical test method, Yekutieli (FDR dependency) at significant level of 0.05 and the Plant GO slim option.

## 3. Results

### 3.1. Phosphorylation of ABI4 by MAPKs

We first set out to check the interaction between ABI4 and MAPKs in our system. Using bimolecular fluorescence complementation we confirmed interactions between ABI4 and both MPK3 and MPK6 in nuclei of *N. benthamiana* leaves transiently expressing these proteins (Supplemental Figure S1). We then used purified recombinant proteins to confirm the *in vitro* phosphorylation of ABI4 by MPK3, MPK4, and MPK6. We chose to use the basal activity of the protein kinases, rather than the enhanced activity following their phosphorylation by MAPKKs, because the basal activity may phosphorylate sites with higher affinity, that cannot be distinguished from lower affinity sites that may be phosphorylated by the enhanced kinase activities attained by the phosphorylation of a MAPK by a MAPKK. Although our data confirmed the reported finding that ABI4 could be phosphorylated by MPK3, MPK4, and MPK6, these kinases varied in their potency to phosphorylate ABI4: all three kinases showed similar degrees of autophosphorylation, but MPK3 was more effective than MPK4 and MPK6 for phosphorylating ABI4 (Figure 1). Thus, we chose to focus on MPK3.

**Figure 1.**
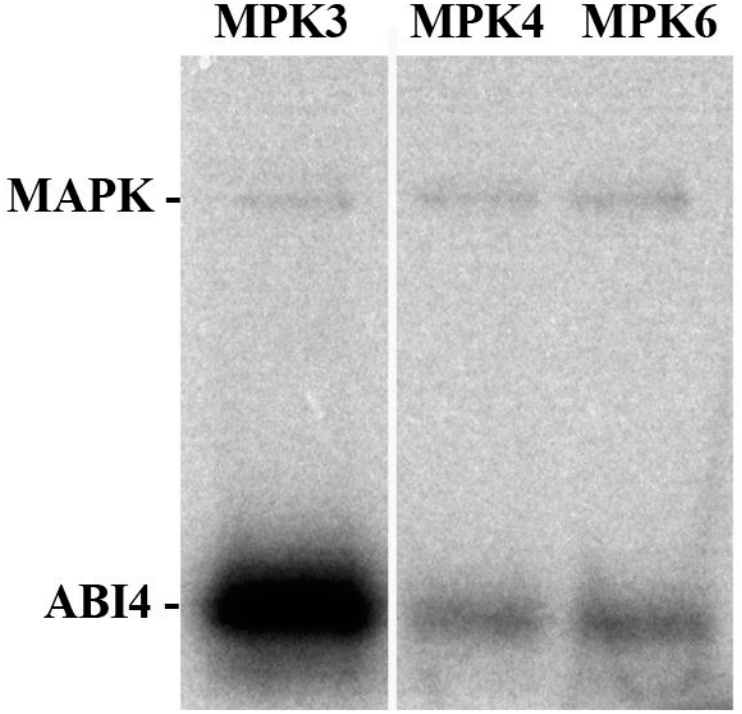
ABI4 phosphorylation by MAPKs. Recombinant ABI4 was incubated with GST-tagged MPK3, MPK4, and MPK6 in the presence of [γ–^32^P]ATP. Samples were resolved on 10% SDS-PAGE, fixed by staining with coomassie-blue, dried, and exposed to a phosphorimager screen. The radiolabeled bands of MAPK and ABI4 are marked. Autophosphorylation of MAPK serves as a control for the kinase activity. The image of the coomassie blue-stained gel is presented in Supplemental Figure S2.

### 3.2. ABI4 Phosphorylation sites

MAPKs phosphorylate serine and threonine residues in their substrate proteins [36]. MAPKs are well studied in eukaryotes, and thus many of their phosphorylation sites are characterized. Two consensus sequences, P-X-S/T-P and S/T-P were found to be highly abundant in protein substrates of MAPKs [37]. Searching the primary amino acid sequence of ABI4 for these motifs, we found that ABI4 has one of the 4-amino acid long MAPK phosphorylation consensus motifs, with S114 as the suggested phosphorylated residue, and two shorter 2-amino acid motifs at residues S130 and T111 (Supplemental Figure S3). Since the short T111 motif overlaps with the longer S114 motif, we first tested whether MPK3 could phosphorylate Ser residues 114 and 130 of ABI4 by mutating the respective codons to encode Ala, a residue that cannot be phosphorylated. Single and double mutations were constructed and verified by DNA sequencing. Wild-type and mutated His-tagged recombinant ABI4 proteins were expressed in *E. coli*, purified by affinity chromatography, and assayed for phosphorylation by MPK3. Replacing S114 of ABI4 with Ala greatly diminished MPK3’s ability to phosphorylate ABI4, whereas mutating S130 did not affect the extent of the observed phosphorylation of ABI4 by MPK3 (Figure 2). The lack of phosphorylation of the S114A/S130A double mutant of ABI4 by MPK3 also suggested that T111 is not a preferred phosphorylation site of this protein kinase.

**Figure 2.**
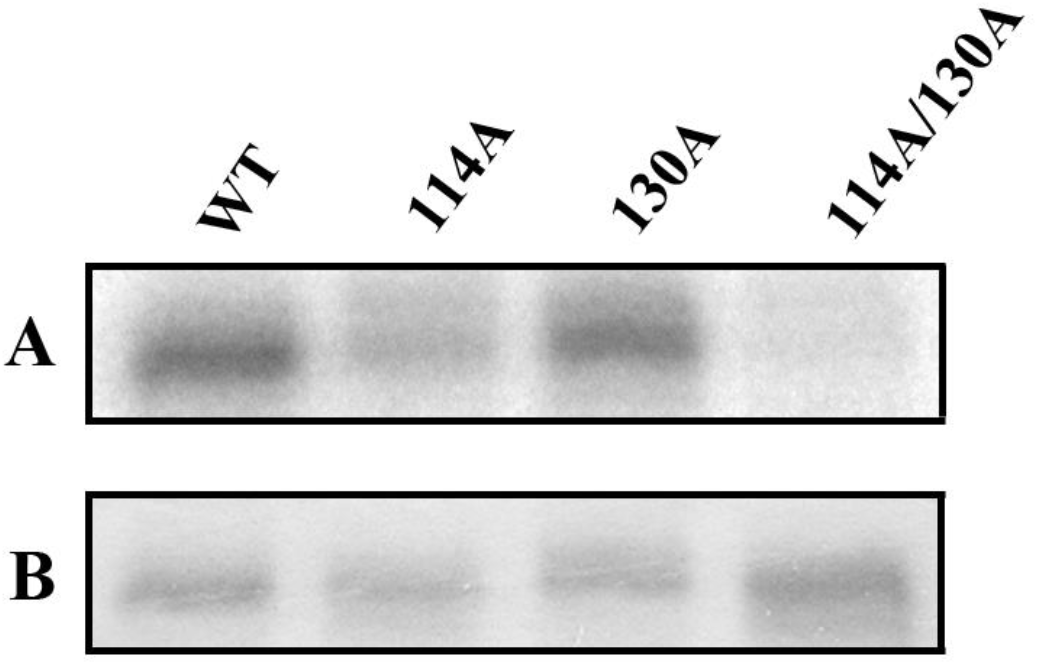
Test of phosphorylation of S114A and S130E mutants of ABI4 by MPK3. The indicated His-tagged ABI4 proteins (WT and the respective Ser to Ala mutants) were expressed in *E. coli*, affinity purified and assayed for phosphorylation by MPK3 in the presence of [γ–^32^P]ATP. Proteins were resolved by SDS-PAGE, gels were stained with coomassie blue, dried, and exposed to a phosphorimager screen. A: Autoradiogram showing phosphorylation of ABI4. B Coomassie staining of the different ABI4 proteins.

### 3.3. Construction of transgenic Arabidopsis plants expressing ABI4 harboring mutations in codons of S114 and S130

To evaluate if the observed *in vitro* phosphorylation of ABI4 by MPK3 (Figure 2) has any biological significance, we tested the potency of ABI4 proteins mutated in S114 and S130 to complement the phenotype of *abi4* mutants. Since overexpression of ABI4 using a strong promoter severely inhibits growth [15, 17], we chose to use the 2 kb ABI4 promoter that was sufficient for the regulated expression of this gene [15]. Codons of ABI4 encoding S114 and S130 were mutated to codons encoding Ala, resulting in phosphorylation-null forms of the protein. A phosphomimetic ABI4 mutant was prepared by replacing the respective Ser residues with Glu, which resembles the negatively charged phosphorylated Ser residue. *Arabidopsis thaliana abi4-1* mutant plants were transformed by *Agrobacteria* carrying pCAMBIA plasmids encoding wild-type ABI4(114S/130S) or the mutated proteins ABI4(S114A), ABI4(S130A), ABI4(S114E), and ABI4(S130E). Homozygous T3 plants resulting from the integration of single T-DNAs were selected by germination and growth in the presence of kanamycin. The phenotype of the resulting transgenic plants was indistinguishable from that of wild-type or *abi4* plants when assayed for germination on plates containing solid MS medium with 0.5% sucrose and 0.55% plant agar, or growth in pots (Supplemental Figure S4).

### 3.4. Germination in the presence of ABA

A pronounced phenotype of *abi4* mutants is their ability to germinate in the presence of inhibitory concentrations of ABA [1]. To test the functional significance of the MAPK phosphorylation sites, seeds of transgenic and control lines were sown on 0.5x MS plates containing 10 μM ABA, and radicle emergence was scored daily for 7 days (Figure 3). All tested genotypes fully germinated with similar kinetics when sown on agar plates containing 0.5x MS medium supplemented with 0.5% sucrose. In agreement with the known phenotype of *abi4*, seeds of the mutant line were less sensitive to ABA inhibition than wild-type plants. For example, 7 days after sowing on plates containing 10 μM ABA, the *abi4* seeds fully germinated whereas only half of the wild type seeds showed radicle emergence. Transformation of *abi4* mutant plants with a construct encoding wild-type ABI4 whose expression was driven by the *ABI4* promoter complemented the germination phenotype (Figure 3, green). Although expression of the ABI4(S114A) phosphorylation-null protein did not complement the germination phenotype, transformants with a construct encoding phosphomimetic ABI4(S114E) yielded seeds with even higher ABA sensitivity than those expressing the wild-type protein. In contrast, the transformation of the *abi4* mutant plants with constructs expressing wild-type, S130A or S130E forms of ABI4 all completely restored the ABA sensitivity. These results clearly suggest that the phosphorylation of S114 but not S130 is necessary for the activity of ABI4.

**Figure 3.**
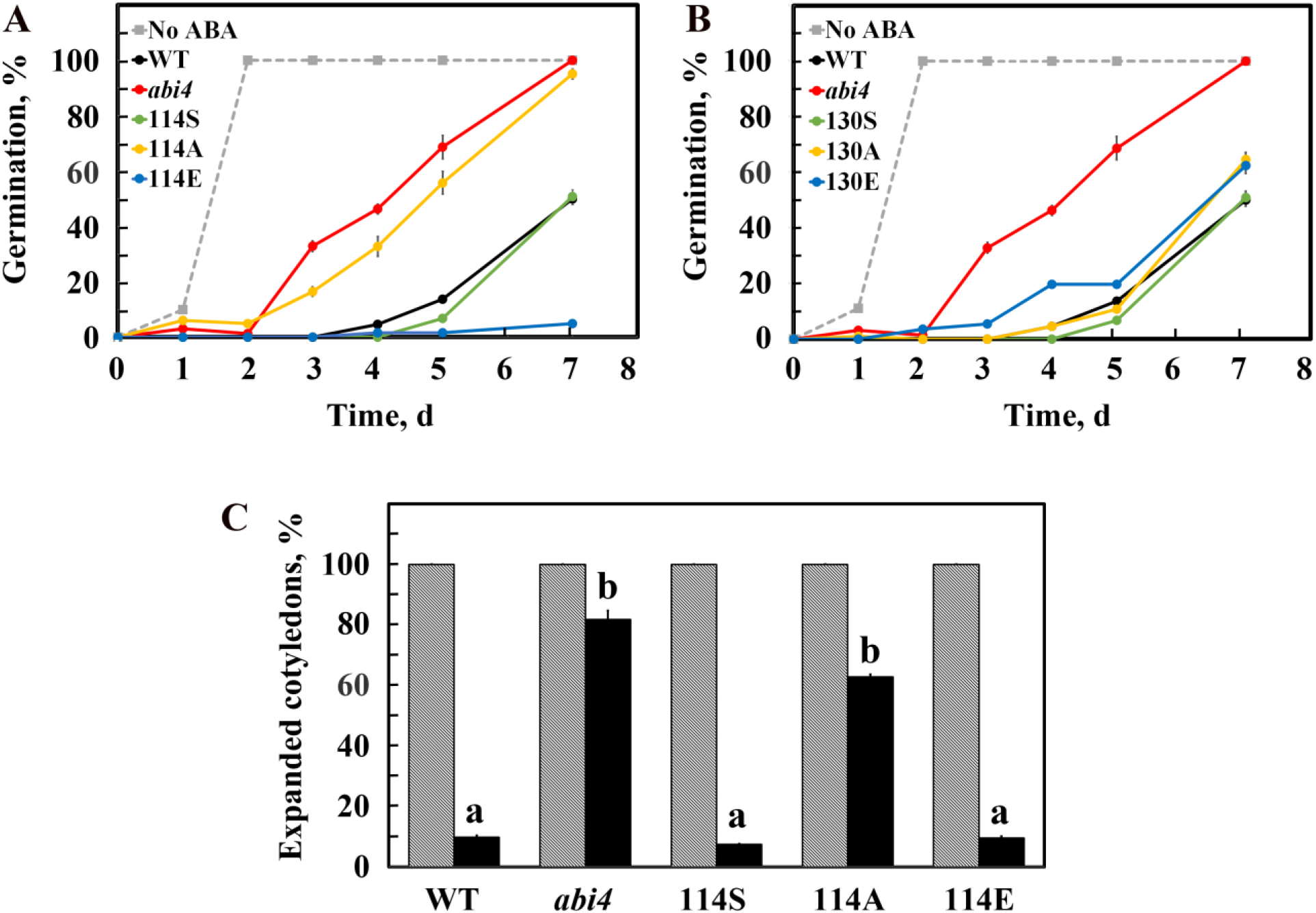
Effect of ABA on seed germination. Surface sterilized cold treated seeds of the indicated genotypes were sown on plates containing agar solidified growth medium without (dashed lines) or supplemented with 10 μM (solid lines) ABA. A&B, Radicle emergence was scored daily. Data represent mean ± SE. A. Mutants of S114. B. Mutants of S130. Dashed gray – germination of all tested genotypes in the absence of ABA. Black, WT plants; Red, *abi4* mutants; Green, Orange and Blue, *abi-4* plants transformed with constructs expressing ABI4 in which amino acid residues 114 (A) or 130 (B) are Ser, Ala or Glu, respectively. C. Expanded cotyledons were scored 10 days after sowing on medium without (shaded) or with (black) 10 μM ABA. The phosphomimetic ABI4(S114E) mutant but not the phosphorylation null ABI4(S114A) mutant could complement the *abi4* phenotype. Data represent mean ± SE. Bars with different letters represent statistically different values using Tukey’s HSD post-hoc test (P ≤0.05).

Following radicle emergence, we also scored the development of cotyledon expansion and greening 10 days after sowing on media supplemented with ABA, in which 9.8% of the wildtype seeds developed green expanded cotyledons (Figure 3C). In contrast, 81.8 % of the *abi4* seeds developed green expanded cotyledons under these conditions. Cotyledon development in the plants expressing ABI4(S114A) was similar to that of the parental *abi4* plants, whereas those expressing ABI4(114S) and ABI4(S114E) resembled those of the wild-type plants, indicating that these constructs express biologically active forms of the ABI4 protein.

### 3.5. Seedling sensitivity to high glucose

High glucose and sucrose concentrations are known to inhibit seed germination and seedling development, and multiple *abi4* mutants have been selected as resistant to these sugars [4–6]. We thus tested seedling development in the presence of 7% glucose. The plants were grown at 22 °C with 16 h light and 8 h dark conditions. Although no differences were obtained in the seed germination rate, the tested genotypes showed differences in cotyledon development rate at 4 days after sowing. Under these conditions, very few seeds of the wild-type plants had expanded cotyledons (Figure 4). In contrast, 89% of the seeds of the *abi4* mutants developed open cotyledons 4 days after plating. The wild-type phenotype could be restored by complementation of the *abi4* mutants with constructs expressing wild-type ABI4(114S) or the phosphomimetic ABI4(S114E) mutant, but not by the phosphorylation-null S114A form of ABI4. Preliminary experiments showed that mutating S130 to Ala or Glu did not affect the protein potency to complement the mutated phenotype, similar to their lack of effects on ABA sensitivity.

**Figure 4.**
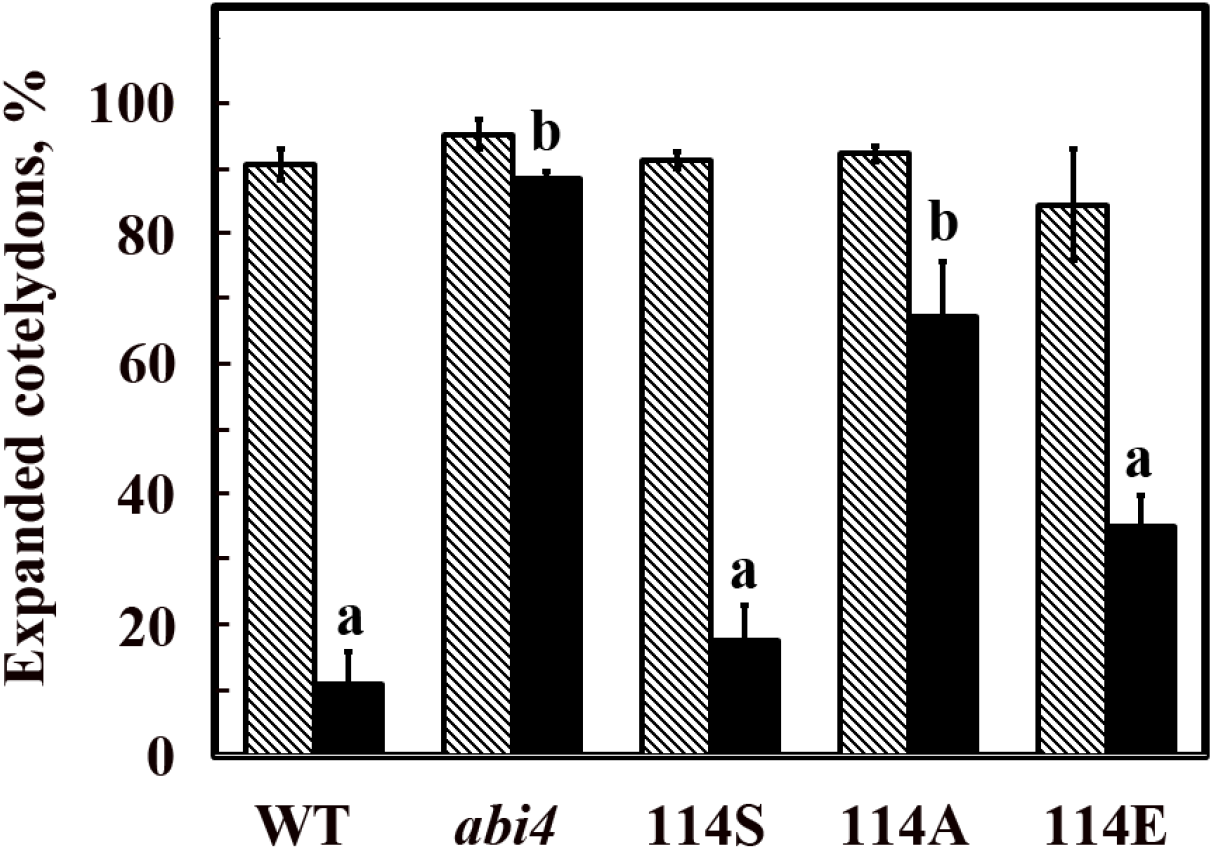
ABI4(S114E), but not ABI4(S114A) mutations, complement *abi4* resistance to glucose effects on cotyledon development. Seeds of plants from the indicated genotypes were sown on plates containing 0.5 x MS salts and 0.7% agar (grey bars) or the same medium supplemented with 7% glucose. Expanded cotyledons were scored 4 days later. Grey and black bars represent data obtained in without or with glucose, respectively. Data represent mean ± SE. Bars with different letters represent statistically different values using Tukey’s HSD post-hoc test (P ≤0.05).

### 3.6. Response to salt

The salt-tolerant *salobreño*(*sañ*) mutants were selected for germination of Arabidopsis seeds in the presence of 250 mM NaCl [3]. One of the isolated mutants, *sañ5*, also termed *abi4-2* resulted from a frameshift mutation in codon 94 of the *ABI4* gene. We thus tested the seeds of *abi4* mutants transformed with constructs expressing the different phosphorylation forms of ABI4 for their potency to germinate in the presence of 0.2 M NaCl. Whereas 92% of the *abi4* seeds germinated 3 days after sowing, only 29% of the wild-type seeds germinated under the same conditions (Figure 5A). Transformation of the *abi4* plants with constructs expressing the wild-type (114S) or phosphomimetic (S114E) form of ABI4 restored sensitivity to salt for inhibition of germination, whereas transformation with the phosphorylation-null S114A form was ineffective, reaching 89% germination at 3 days.

**Figure 5.**
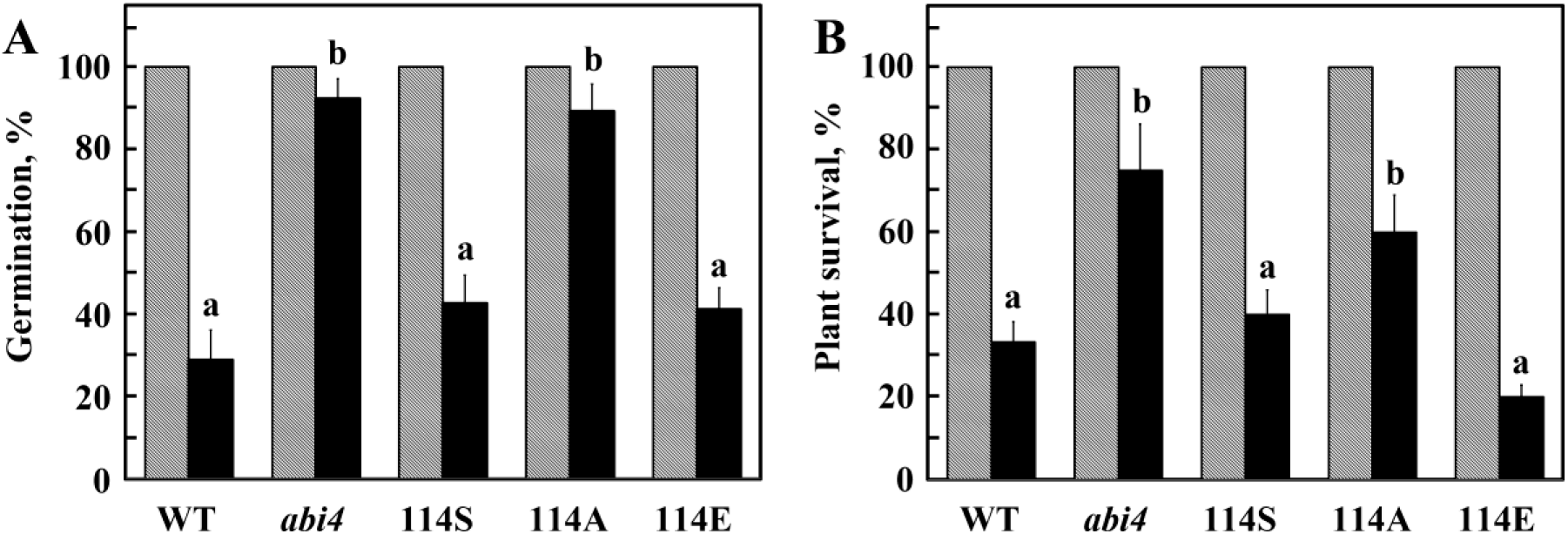
ABI4(S114E), but not ABI4(S114A) mutations, complement *abi4* resistance to NaCl effects on seed germination and viability of rosettes. A. Seeds from the indicated genotypes were sown on plates containing 0.5 x MS salts, 0.5% sucrose and 0.7% agar, containing in addition 0 (shaded) or 0.2 M NaCl (black). Radicle emergence was scored 3 days later. Data represent mean ± SE. Bars with different letters represent statistically different values using Tukey’s HSD post-hoc test (P≤0.01). B. Seeds of the indicated genotypes were sown in a pot containing potting mix and irrigated with water for one month. Rosettes were than irrigated twice a week with water (shaded) or 0.25 M NaCl and 16 mM CaCl_2_. Plant survival was scored 17 d later. Data represent mean ± SE. Bars with different letters represent statistically different values using Tukey’s HSD post-hoc test (P≤0.01).

We previously showed that *abi4* mutants display higher salt tolerance when exposed to NaCl at the rosette stage [26]. To test the relevance of ABI4 phosphorylation for this response, one-month-old pot grown plants were irrigated with 0.25 M NaCl and 16 mM CaCl_2_, and plant survival was scored 17 d later. Na^+^/Ca^2+^ ratios of 10-20 were suggested to protect cell membranes [38]. In agreement with earlier studies, *abi4* mutants displayed enhanced salt tolerance compared with wild-type plants (Figure 5). This tolerance was reversed by complementation with wild-type ABI4(114S) and ABI4(S114E) phosphomimetic mutant proteins, but not by a transgene encoding the phosphorylation-null ABI4(S114A) mutant.

### 3.7. Lateral root development

The *A. thaliana abi4* mutants produce more lateral roots than wild-type plants, whereas ABI4 expression inhibits lateral root development [15, 39]. Seeds were sown in Petri dishes containing 0.5x MS, 0.5% sucrose and 0.5% plant agar, and plates were incubated for 12 days in a growth room in 12-hour light/dark conditions. The plants were harvested, and the lengths of the primary roots (PR) and first lateral roots (LR) were measured. The lengths of the PR did not differ among genotypes, whereas the length of the LR and the calculated ratios of the LR/PR lengths of the *abi4* mutant, and plants expressing ABI4(S114A) were significantly higher than those of the wild-type, and plants expressing ABI4(114S) and ABI4(S114E) (Figure 6).

**Figure 6.**
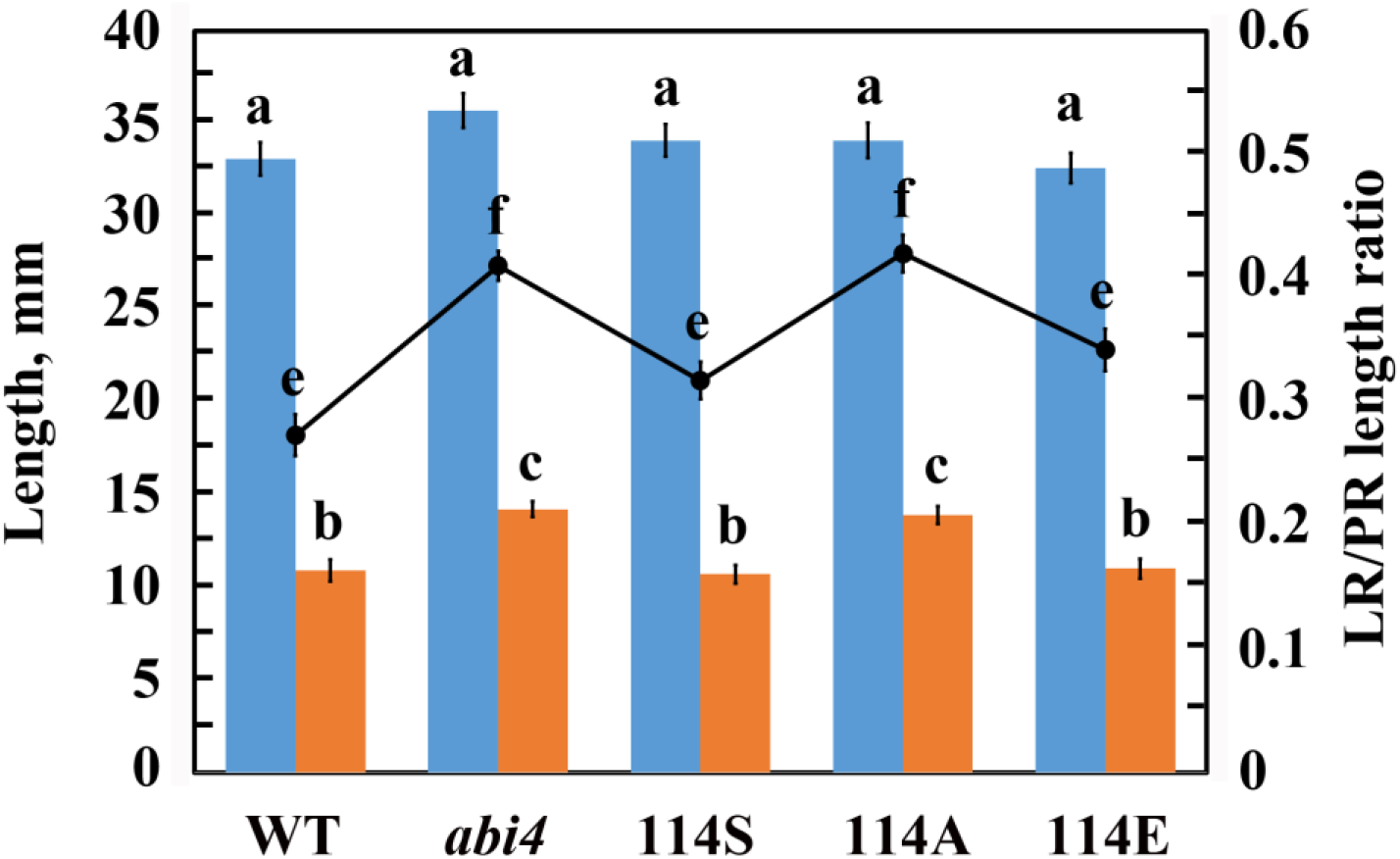
ABI4(S114E), but not ABI4(S114A) mutations, complement over-production of lateral roots by *abi4* mutants. Length of primary (blue bars) and 1^st^ lateral roots (orange bars) in 12 d old seedlings were measured. (●) The ratio of lateral root/primary root lengths. Data represent mean ± SE. Bars with different letters represent statistically different values using Tukey’s HSD post-hoc test (P ≤0.01).

### 3.8. Phosphorylation of Serine 114 of ABI4 is essential for modulating gene expression in planta

ABI4 is known to act as both an activator and a repressor of transcription. In previous studies we shown that ABI4 is expressed in roots and modulates root function [15, 26]. To determine whether the phosphorylation state of the S114 residue affects transcriptional control by ABI4, we analyzed the root transcriptomes of 1-week-old seedlings of the 5 genotypes following a 4 hr treatment with 50 μM ABA. Principal component analyses show that wild-type and *abi4* transcriptomes display the greatest difference in this set, with all 3 transgenic lines intermediate with respect to PCA1 (Supplemental Figure S5).

We first compared the transcriptomes of ABA-treated roots of wild-type and *abi4* seedlings to identify the complete set of ABI4-regulated genes (see Methods). This analysis identified 823 genes, differentially expressed between the two genotypes, where the expression levels of 592 and 231 genes were higher and lower, respectively, in ABA-treated roots of the *abi4* mutant (Supplemental Figure S6, and Supplemental Table S2). Our results also showed that transformation of *abi4* mutants with a construct expressing wild-type ABI4(114S) protein, partially restored the transcription pattern of the wild-type plants. Nevertheless, as expressing ABI4(114S) in *abi4* mutants was sufficient for complementation of the lateral root development phenotype (Figure 6), as well as ABA and stress responses at germination and seedling growth of the *abi4* mutant (Figures 3-5), we focused our analysis on 420 genes whose expression is significantly different between roots of *abi4* mutants and both wild-type and *abi4* plants expressing wild type ABI4(114S) (Figure 7A and Supplemental Table S3). Of these, the expression levels of 355 genes (84.5%) were higher in *abi4* mutant roots, suggesting that ABI4 acts mostly as a repressor.

**Figure 7.**
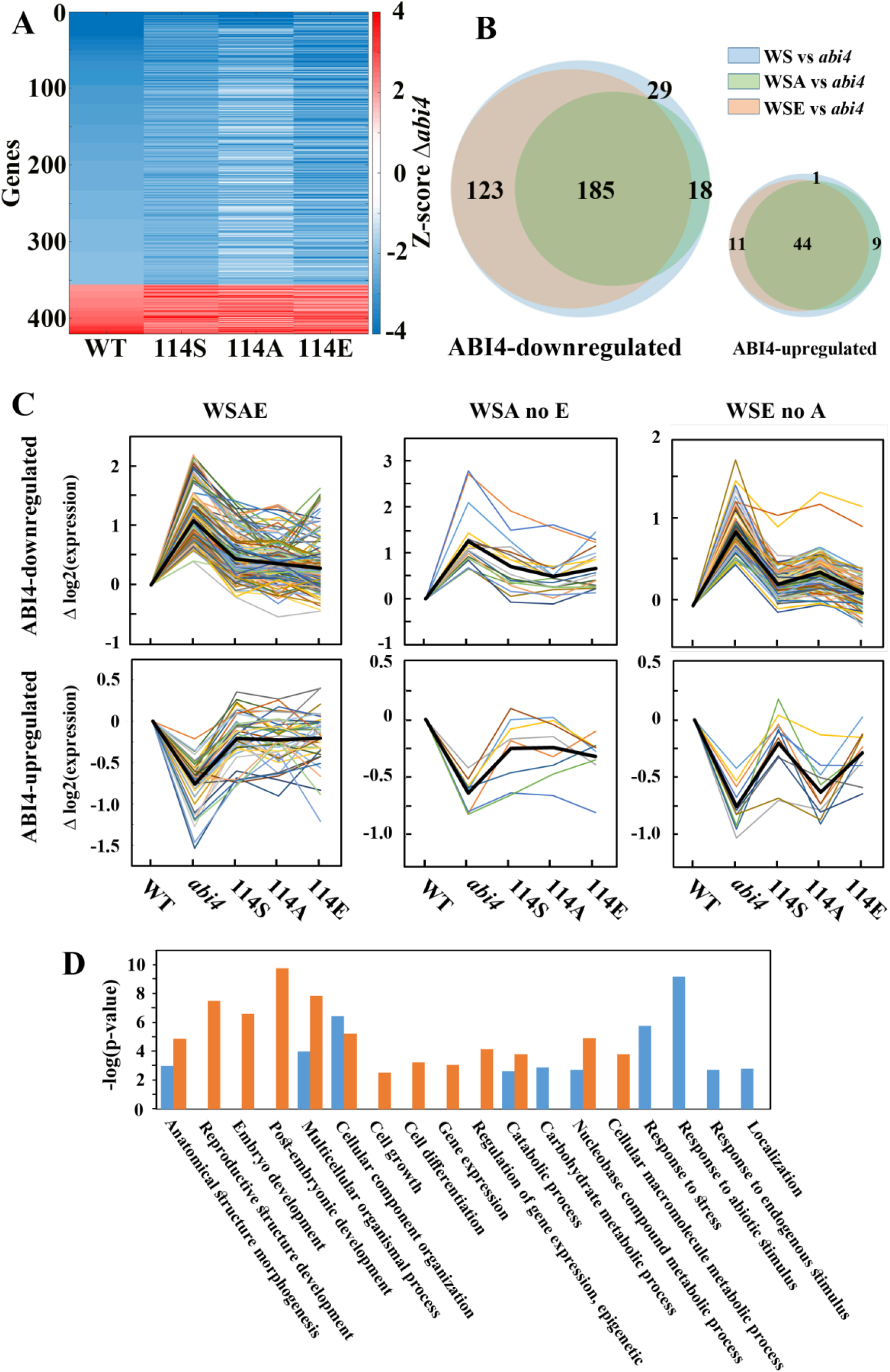
Expression of ABI4-regulated genes that are affected by the phosphorylation status of S114. A. Heatmap of genes that are differentially expressed both in the wild-type and *abi4* complemented by ABI4(114S) genotypes versus *abi4.* Color indicate the Z-score values of the differential expression of the indicated genotypes compared to *abi4* (see Methods). B. Venn diagram presenting ABI4-regulated genes shown in A. WS, genes that are differentially expressed both in wild type and ABI4(114S); WSA, genes that are differentially expressed in WS and ABI4(S114A); WSE, WS and ABI4(S114E). C. Expression patterns of gene clusters whose expression is independent of the phosphorylation state of ABI4-S114 (WSAE), or dependent on the non-phosphorylated (WSA no E) or phosphorylated (WSE no A) states of ABI4 S114, corresponding to the Figure 7B. Plots show log2 values of medians of each gene in the group normalized to the expression of the respective gene in the wild-type plant, where plots of individual genes are shown by colored lines, and the average values in thick black lines. D. Biological processes of the proteins encoded by the ABI4-downregulated genes in (B). Biological processes were analyzed by AgriGo ver2. Blue bars represent ABI4-down regulated genes whose expression is not affected by the phosphorylation state of ABI4 (WSAE); orange bars represent genes in group WSE, that are down-regulated by the phosphorylated state of S114 of ABI4 (WSE no A). The WSA no E group was too small for the analysis.

To determine whether phosphorylation state affected ABI4 activity *in planta*, we compared the expression levels of these genes in the transgenic plants expressing ABI4(114S), ABI4(S114A) and ABI4(S114E) relative to the respective levels in the *abi4* plants to determine whether phosphorylation state affected ABI4 activity *in planta* (Figure 7A and Supplemental Figure S7). Heat map and Venn diagrams show that the gene expression profile in roots of ABA-treated plants expressing the phosphomimetic ABI4(S114E) protein more closely resembles that of the wild-type ABI4(114S) profile than that of plants expressing the phosphorylation-null ABI4(S114A) (Figure 7A & B). Expression levels of 52% and 68% of the genes whose transcript levels were higher or lower, respectively, in *abi4* mutants than in wild-type and *abi4* expressing ABI4(114S) plants, were reversed by either ABI4(S114A) or ABI4(S114E) proteins (Figure 7B & C, marked as WSAE), suggesting that modulation of the expression of these genes by ABI4 is independent of the phosphorylation state of S114. In contrast, the phosphorylated state of S114 is required for modulation of the expression of 35% and 17% of the ABI4-regulated genes whose levels in *abi4* were higher or lower, respectively, than in *abi4* mutants expressing ABI4(S114E), but not ABI4(S114A) (Figure 7B & C, marked as WSE no A). Even fewer ABI4-modulated genes require the non-phosphorylated form of S114 for regulation: only 5% and 14% for repression or activation, respectively (Figure 7B and C, marked as WSA no E). The list of genes affected by phosphorylation state is presented in Supplemental Tables S4 – S6.

The genes that require phosphorylation of S114 for repression are enriched for those affecting development (embryo, post-embryonic and reproductive), morphogenesis, cell growth, differentiation, and regulation of gene expression. Those that are repressed regardless of phosphorylation state are enriched for responses to stress, abiotic and endogenous stimuli (Figure 7D). The few ABI4-induced genes that require phosphorylation of S114 include several that are involved in defense response and/or are induced by salt treatment of roots [40].

In terms of molecular function, the largest classes of genes whose expression is modulated by the phosphorylation of S114 encode nucleic acid binding proteins and enzymes (Supplemental Figure S8). Consistent with the ABA-resistant growth of the *abi4* mutants, the ABI4-repressed genes that were most highly enriched relative to their abundance in the genome code for factors needed for growth including nucleic acid binding proteins mediating transcription, RNA splicing, chromatin modification; translation initiation and elongation factors; and ion channels.

Screening the upstream sequences of the differentially expressed genes for 6 DNA sequences previously reported to be bound by ABI4 identified an average of 2.55 putative ABI4 binding sites per gene. The most abundant are the CE1-like element (CACCK), ABRE-like sequence (BACGTGKM), and DRE core motif (RCCGAC) (Supplemental Figure S9). The differences between the numbers of the different binding sites in genes whose expression levels is affected by the different phosphorylation states of S114 of ABI4 maybe too small to be significant. Moreover, these sites are present in similar abundance in all the root expressed genes (Supplemental Figure S9), suggesting that the occurrence of these sites in the promoter DNA sequence is not enough to be regulated by ABI4. Furthermore, some of these genes may be indirectly regulated by ABI4 so would not contain ABI4 binding sites in their promoters.

## 4. Discussion

ABI4 activity has been shown to be regulated at the level of gene expression, protein stability, and most recently by phosphorylation. Previous studies of ABI4 phosphorylation by MAP kinases identified 3 MAPKs that could phosphorylate ABI4 *in vitro* [20] and focused on how MAPKs activated by MAPKKs affected ABI4 function in chloroplast retrograde signaling [24]. Guo et al. [24] identified 3 residues phosphorylated by MPK3 and MPK6: T111, S114, and S130. Although the relevance of each phosphosite was not tested individually, a mutant form of ABI4 with all 3 residues converted to Ala was unable to bind an *LHCB* promoter *in vitro* or repress *LHCB* expression in transgenic plants. More recently, phosphorylation of ABI4 by MPK3 and MPK6 was found to repress production of adventitious roots at the root-hypocotyl junction, in part by inhibiting accumulation of reactive oxygen species and programmed cell death in overlying cell layers [25]. Furthermore, these MPKs phosphorylate PP2C12, which dephosphorylates ABI4, thereby inactivating PP2C12 and stabilizing ABI4 in an activated state. As for *LHCB* regulation, this study did not distinguish functional significance among the 3 potential phosphosites.

In the current study, we focused on activation of ABI4 by MPK3, the kinase with the highest intrinsic affinity for ABI4 without prior activation by a MAPKK. Under these conditions, substitution of both S114 and S130 with Ala was necessary and sufficient to block phosphorylation of ABI4, suggesting that T111 is not phosphorylated by MPK3 displaying its basal activity (Figure 2). Furthermore, the extent of phosphorylation suggests that the phosphorylation of S114 is preferred over S130 (Figure 2). Although both S114 and S130 were phosphorylated *in vitro*, functional tests of the individual phosphosites showed that phosphorylation of S114 was necessary for ABI4 activity in promoting inhibition of growth by ABA, high sugar or salinity, including inhibition of lateral root growth. MPK3 was reported to be involved in a number of plant stresses [22], consistant with a possible physiological role in the modulation of ABI4 activity: MAPK3 activity was enhanced by ABA [41] and NaCl [42]. In addition, although there is very little information on the role of MAPKs in sugar signaling, MAPK6 was demonstrated to be involved in ABA and sugar signaling in seed germination in Arabidopsis [43], suggesting that other MPKs may also be involved.

The root transcriptome of the ABI4(S114A) plant is most similar to that of the *abi4* mutant, whereas the transcriptomes of ABI4(S114) and ABI4(S114E) are closer to that of wild type plants. It is not clear why transgenic expression of the wild type ABI4 protein restored regulation of only half of the genes mis-regulated in the *abi4* mutant, but this might reflect different threshold levels of ABI4 needed for proper expression of its targets. If so, even small differences in expression or accumulation of ABI4 could account for this incomplete complementation, and would be consistent with the transcriptome variability observed among biological replicates of any given genotype.

ABI4-regulated individual genes and transcriptomes have been reported in a variety of contexts, including seed development, dormancy and germination [12, 13, 44], oxidative stress response [45], ABA-treatment of ectopic expression lines [34], salt stress response [46], and retrograde signaling from organelles [9, 47, 48]. In dry seeds, expression of over 2200 genes was reduced at least two-fold in *abi4* mutants, while over 2800 genes were expressed at least twofold higher in this background [12], reflecting primarily repressive action of ABI4. Approximately 50-70% of genes activated by ABA treatment of ABI4 overexpression lines were also activated in wild-type plants exposed to ABA, high salt or osmotic stress at different ages [34]. In contrast, consistent with the diversity of physiological and developmental contexts described above, there is very little overlap between the ABI4 target genes identified in these studies. A further complication for making comparisons is that many ABA or stress responses proceed through phases of early vs. late gene expression, so different studies may simply be examining different phases or transient responses [49]. However, all of these studies show that ABI4 can function as either an activator or repressor of distinct subsets of genes.

The present study analyzed ABI4-regulated genes in seedling roots following a brief exposure to ABA. Comparison of wild-type and *abi4* mutants identified at least 2.5-fold more genes that are repressed by ABI4 under these conditions than genes that are activated (Supplemental Figure S6). Filtering out those genes whose expression was not restored to wildtype levels by the complementing transgene removed about half of these genes, but preferentially removed ABI4-induced genes, resulting in a 4-fold difference between repressed and induced genes. Further narrowing the focus to genes whose regulation was restored in roots of plants expressing ABI4(S114E) but not in ABI4(S114A) seedling roots, the ratio of repressed to induced genes was greater than 10-fold, suggesting that phosphorylation of S114 of ABI4 might be more important for action as a repressor. Whereas the genes requiring phosphorylation of ABI4 for repression were enriched in functions associated with reproductive development and growth, most of the few transcripts that were higher in the ABA-treated roots of wild-type and ABI4(S114E) than *abi4* were associated with abiotic and/or biotic stress responses.

Many transcription factors are known to require phosphorylation for activity. For example, in the core ABA signaling pathway, stability and/or activity of the ABA and stress-responsive ABI5/ABF/AREB clade of bZIPs depend on phosphorylation by SnRK2 family protein kinases [50], possibly mediated by interactions with 14-3-3 proteins [51]. However, the effects of phosphorylation of these transcription factors on accumulation of specific transcripts has not been systematically analyzed. Our results show that phosphorylation of a single residue of ABI4 affects its ability to regulate some, but not all, ABI4 target genes. Similar results were described for the neurogenic transcription factor, SOX11, in transfected cells [52]. In this case, transactivation assays testing wild-type, phosphomimetic and phosphorylation-null mutants for effects on three different promoter-LUC fusions produced three distinct results: no difference between wild type and mutant SOX11, and both phosphomutants more active than wild type but differing in whether phosphomimetic or phosphorylation-null was more effective. These results suggest that the balance between phosphorylation states may be another important aspect regulating activity relative to specific target genes.

Surprisingly, some of the genes that are repressed in ABA-treated roots by ABI4, regardless of its phosphorylation state, show reduced expression in *abi4* mutant seeds (e.g., AT3G53040, encoding a LEA protein) [12], indicating that ABI4 may be either an activator or repressor of a given gene in different developmental contexts. This may reflect interactions with a variety of other transcription factors. The putative ABI4 binding sites have diverse sequences and previous EMSA studies indicate relatively low affinity and specificity of binding to several of these [26, 34], suggesting that *in vivo* binding may be modulated by associations with other factors. Two recent reports document formation of complexes containing ABI4 and the histone deacetylase HDA9, leading to repression of a subset of ABI4-regulated genes by chromatin modification during drought stress [53, 54]. Furthermore, a recent report indicates that intrinsically disordered domains in transcription factors can influence *in vivo* binding site specificity [55], and more than half of ABI4 is predicted to be disordered.

In summary, our results confirm previously reported MAPK phosphorylation sites, but show that phosphorylation of only one of these is actually necessary for ABI4 activity in responding to ABA, glucose and salt stress. Furthermore, the transcriptome experiment identified a novel subset of ABI4-regulated genes, approximately one-third of which are dependent on phosphorylation of ABI4 for proper expression in roots, and this regulation is primarily repressive.

## Accession Numbers

The SRA accession number for the RNA-seq data reported in this paper is RJNA655114. Arabidopsis genes studied: ABI4, At2G40220; MPK3, At3G45640; MPK4, At4G01370; MPK6, At2G43790.

## Author contribution statement

N.E. performed most of the experiments and participated in writing the draft of the manuscript; T.M. performed some experiments; E.C.S. cloned, expressed and purified the recombinant MAPKs; Da.B.Z. and S.B. analyzed the RNA seq data and participate in writing the manuscript; R.F. contributed to experiment planning, data analysis and writing the manuscript; Du.B.Z. conceived the project idea, designed and analyzed the experiments, wrote part of the manuscript.

## Funding

This work was supported by a grant No. 2011097 from the US Israel Binational Science Foundation (BSF) (to R.F. and Du.B.Z.).

## Declaration of Competing Interest

The authors declare that there is no known competing interest in this submission

## Acknowledgments

Du.B.Z. is the incumbent of The Israel and Bernard Nichunsky Chair in Desert Agriculture, Ben-Gurion University of the Negev

## Appendix A. Supplementary data

**Supplemental Figure S1.** BiFC interactions between ABI4 and MPK3 and MPK6.

**Supplemental Figure S2.** Coomassie blue-stained gel of ABI4 phosphorylation by MAPKs.

**Supplemental Figure S3**. Putative MAPK phosphorylation sites in ABI4.

**Supplemental Figure S4.** Phenotype of the studies genotype grown in standard conditions.

**Supplemental Figure S5**. Principal component (PCA) analysis of gene expression levels in roots of ABA-treated plants.

**Supplemental Figure S6.** Expression of ABI4-regulated genes in roots of ABA-treated plants.

**Supplemental Figure S7.** Heatmap of genes that are differentially expressed both in the wildtype and abi4 complemented by ABI4(114S) genotypes versus abi4.

**Supplemental Figure S8.** Molecular function of the differential expressed genes.

**Supplemental Figure S9.** Analysis of putative ABI4-binding sites in the differentially expressed genes.

**Supplemental Table S1.** List of primers used in this study.

**Supplemental Table S2.** List of genes with different root expression between ABA-treated wildtype and *abi4* mutant plants.

**Supplemental Table S3.** List of differentially expressed genes intersecting between the transgenic plant expressing ABI4(114S), ABI4(S114A) and ABI4(S114E).

**Supplemental Table S4.** List of genes with different root expression in both wild-type and 114S compared to abi4, that are also differentially expressed (|z-score|>2) in both 114A and 114E (WSAE).

**Supplemental Table S5.** List of genes with different root expression in both wild-type and 114S compared to abi4, that are also differentially expressed (|z-score|>2) in 114E but not in 114A (WSE no A).

**Supplemental Table S6.** List of genes with different root expression in both wild-type and 114S compared to abi4, that are also differentially expressed (|z-score|>2) in 114A but not in 114E (WSA no E)

## Notes

### Competing Interest Statement

The authors have declared no competing interest.

### Summary of Updates

Manuscript Text, Figure 3 and Supplemental files revised, Authors affiliations updated.

